# DNA polymerase β prevents AID-instigated mutagenic non-canonical mismatch DNA repair

**DOI:** 10.1101/2020.01.30.926964

**Authors:** Mahnoush Bahjat, Maria Stratigopoulou, Bas Pilzecker, Tijmen P. van Dam, Simon Mobach, Richard J. Bende, Carel J.M. van Noesel, Heinz Jacobs, Jeroen E.J. Guikema

**Author notes:** These authors contributed equally (Maria Stratigopoulou, Bas Pilzecker). **Correspondence**: Jeroen E.J. Guikema, Ph.D. Amsterdam UMC, Location AMC, University of Amsterdam, department of Pathology, Meibergdreef 9, 1105 AZ, Amsterdam, The Netherlands.

## Abstract

In B cells, the error-prone repair of activation-induced cytidine deaminase (AID)-induced lesions in immunoglobulin variable genes cause somatic hypermutation (SHM) of antibody genes. Due to clonal selection in the germinal centers (GC) this active mutation process provides the molecular basis for antibody affinity maturation. AID deaminates cytosine (C) to create uracil (U) in DNA. Typically, the short patch base excision repair (spBER) effectively restores genomic U lesions. We here demonstrate that GC B cells actively degrade DNA polymerase β (Polβ), resulting in the inactivation of the gap-filling step of spBER. Consequently, lesions instigated by AID, and likely other base damages, are channeled towards mutagenic non-canonical mismatch repair (mncMMR), responsible for the vast majority of mutations at adenine and thymine (A:T) bases. Apparently, GC B cells prohibit faithful spBER, thereby favoring A:T mutagenesis during SHM. Lastly, our data suggest that the loss of Polβ relates to hypoxia that characterizes the GC microenvironment.

## INTRODUCTION

Humoral or antibody-mediated immunity is essential in protecting the host against foreign pathogens. A key feature of B cells in the humoral immune response is their ability to undergo somatic hypermutation (SHM) and immunoglobulin (Ig) class switch recombination (CSR), yielding antibodies with consecutively greater affinity, and of different Ig isotypes (Methot and Di Noia, 2017). These processes occur in the germinal center (GC), specialized anatomical sites that arise within B-cell follicles in the lymphoid organs upon infection or immunization. GCs contain two zones: a dark zone (DZ) and a light zone (LZ). In the DZ, large centroblasts clonally expand by rapid proliferation and undergo SHM. Subsequently, they migrate to the LZ and transit into centrocytes that are subject to clonal selection (MacLennan, 1994). Affinity maturation is based on active mutagenesis of rearranged *Ig* variable V(D)J genes by SHM in centroblasts and subsequent selection of centrocyte subclones expressing higher affinity antibody variants for the cognate antigen and their differentiation into memory B cells and plasma cells (Victora et al., 2010).

During SHM, predominantly single nucleotide substitutions are introduced in −3 the *Ig V* genes at a rate of approximately 10^-3^ mutations per base pair (bp) per generation (McKean et al., 1984; Rajewsky et al., 1987). Activation-induced cytidine deaminase (AID) is highly expressed in GC B cells in the DZ and is of crucial importance for both SHM and CSR (Muramatsu et al., 2000; Revy et al., 2000). AID catalyzes the deamination of cytosine (C) into uracil (U), resulting in the formation of highly mutagenic dU lesions in the DNA. Typically, base damages including deaminations are efficiently repaired in an error-free fashion by base excision repair (BER). However, in GC B cells dU lesions resulting from AID activity are not faithfully restored. Two principal scenarios may explain these mutagenic properties, (i) the active generation of AID-instigated lesions outcompetes canonical, non-mutagenic short patch BER (spBER), or (ii) spBER itself is suppressed to favor non-canonical, mutagenic repair pathways.

The preferred target of AID is the C embedded in the ‘hotspot’ WR<u>C</u>Y motif (RGYW on the opposite strand; W = A/T, R = A/G, Y = C/T) (Álvarez-Prado et al., 2018; Rogozin and Kolchanov, 1992). Processing of the dUs results in base substitutions and/or DNA double strand breaks (DSBs), which is largely determined by the ensuing DNA repair activities (Bahjat and Guikema, 2017; Peled et al., 2008; Stavnezer et al., 2008). AID activity appears to be restricted to the G1-phase of the cell cycle (Le and Maizels, 2015; Schrader et al., 2007; Wang et al., 2017), but dUs not repaired in the G1 phase will instruct a template T during S-phase, leading to insertion of an A on the opposite strand, resulting in C to T and G to A transitions. Alternatively, U removal by uracil-DNA-glycosylase (Ung), a component of the BER pathway, followed by replication over the apurinic/apyrimidinic (AP) site by translesion synthesis (TLS) DNA polymerases such as Rev1 results in transversions at C:G bps (Di Noia and Neuberger, 2002; Krijger et al., 2013; Rada et al., 2002). In addition, AID-dependent cytosine deaminations lead to mutations at A:T bps during SHM, which is based on a mutagenic non-canonical type of MMR (mncMMR). Here, the the dU lesion is recognized as a U:G mismatch by MutS protein homolog 2 (Msh2) and MutS homolog 6 (Msh6) heterodimer, prompting DNA patch excision by the exonuclease 1 (Exo1). Subsequent gap-filling by the error-prone TLS DNA polymerase η (Polη) contributes to A:T mutagenesis, focused on the WA motif (Bardwell et al., 2004; Delbos et al., 2007; Martin et al., 2003; Martomo et al., 2004; Rogozin et al., 2001; Wilson et al., 2005; Zeng et al., 2001). In addition, approximately half of the G:C transversions rely on both Ung and mncMMR (Krijger et al., 2009, 2013). Exo1 requires an incision for entry to initiate patch excision. MutL-alpha (MutLα), composed of MutL homolog 1 (Mlh1) and Pms1 homolog 2 (Pms2), can nick the DNA on the 5’ side of the mismatch via Pms2 endonuclease activity, but Pms2 endonuclease-deficient mice displayed normal A:T mutagenesis (van Oers et al., 2010, 2). However, loss of both Pms2 and Ung resulted in a 50% reduction of A:T mutations, suggesting that both Pms2 and Ung contribute to DNA incision required for Exo1 entry (Girelli Zubani et al., 2017).

In GC B cells, the AP site endonuclease 2 (Ape2) was shown to be primarily responsible for DNA nicking activity downstream of Ung, thereby facilitating CSR and A:T mutagenesis during SHM (Guikema et al., 2007; Stavnezer et al., 2014). AP endonuclease nicking yields a 3’ hydroxyl group and a transient 5’ abasic deoxyribose phosphate (dRP) moiety (Springgate and Liu, 1980). During BER, AP sites are faithfully repaired by Polβ, inserting the correct nucleotide at the site of the nick and removing the dRP via its associated lyase activity (Matsumoto and Kim, 1995). DNA ligase III (LigIII) along with its cofactor x-ray repair cross complementing 1 (Xrcc1) catalyzes the nick-sealing step (Caldecott et al., 1994, 1996; Gao et al., 2011; Simsek et al., 2011). The exact role of Polβ in SHM remains to be established, as *ex vivo* analysis of B cells derived from Polβ-deficient hematopoietic stem cells (HSCs) in reconstituted mice indicated normal SHM (Esposito et al., 2000), whereas *in vitro* stimulated Polβ-deficient B cells showed a small but significant increase in CSR and SHM (Wu and Stavnezer, 2007).

Of interest, *in vitro* systems to study AID-instigated mutagenesis are characterized by the low frequency of A:T mutations versus *in vivo* B cells (approximately 0-20% versus 50-60% of all mutations, respectively) (Martin et al., 2002; Matthews et al., 2014; Xiao et al., 2007; Yoshikawa et al., 2002). We hypothesized that this apparent discrepancy is caused by distinct DNA repair activities *in vitro* versus *in vivo.*

Herein, we show that cells cultured *in vitro* functionally express Polβ, whereas it is expressed at very low levels in GC B cells *in vivo.* We provide evidence that Polβ suppresses AID-instigated A:T mutagenesis *in vitro,* demonstrating that active loss of Polβ skews repair of AID-induced lesions towards mncMMR. Furthermore, our data suggest that GC hypoxia is responsible for the systemic breakdown of Polβ protein.

## RESULTS AND DISCUSSION

### Functional knockdown of Polb enables A:T mutagenesis in CH12-F3 cells

The mouse B lymphoma cell line CH12-F3 can be induced to express AID and undergo CSR to IgA when stimulated with anti-<u>C</u>D40, <u>I</u>nterleukin-4 and <u>T</u>ransforming growth factor β (TGFβ) (CIT) (Muramatsu et al., 1999; Nakamura et al., 1996). In addition, these cells accrue AID-dependent point mutations in the *Ig* switch μ region (Sμ) upon stimulation, similar to mouse splenic B cells (Nagaoka et al., 2002; Schrader et al., 2003). Strikingly, despite a high frequency of AID-dependent mutations 5’ of the Sμ repetitive region, the CH12-F3 cells conspicuously lack A:T mutations for hitherto unknown reasons (Matthews et al., 2014). We hypothesized that the late gap-filling and repair activities of Polβ and LigIII diminishes entry points for Exo1, thereby suppressing A:T mutagenesis. To address this, we used short hairpin RNA (shRNA) interference to silence *Polb* gene expression in CH12-F3 cells (**Fig. S1A**). Functional *Polb* knockdown was confirmed by demonstrating increased cellular sensitivity to the base methylating agent methylmethane sulfonate (MMS), which requires Polβ for DNA repair (Sobol et al., 1996) (**Fig. S1B**). To determine the contribution of Polβ to the resolution of AID-induced lesions, genomic DNA was isolated from CH12-F3 cells after 7 days of CIT stimulation. The region 5’ of the Sμ tandem repeats was PCR amplified from the total cell population and mutations were analyzed by cloning and Sanger sequencing. The overall mutation frequency did not differ significantly between wild type parental (WT) and *Polb* knockdown CH12-F3 cells (2.1 x10^-3^ vs 2.5 x10^-3^, respectively, **Table I**). We found that mutations at A:T base pairs were infrequent in the WT CH12-F3 cells (9 out of 59363 sequenced A:T bps were mutated), in agreement with a previous study (Matthews et al., 2014). In contrast, *Polb* knockdown CH12-F3 cells showed a considerable number of A:T mutations (23 out of 57988 sequenced A:T bps were mutated. We conclude that silencing of *Polb* results in an increase in A:T mutagenesis in CH12-F3 cells (1.4 ×10^-4^ vs 3.9 x10^-4^, *p* = 0.018; 3-fold increase, **Fig. 1**). We did not observe any significant differences in the frequency of G:C transitions (4.2 ×10^-3^ vs 5.0 ×10^-3^), G:C transversions (2.7 ×10^-4^ vs 2.4 ×10^-4^) and AID hotspot mutations (4 ×10^-3^ vs 5 x10^-3^, **Table I**, **Fig. 1**). Based on these results we concluded that *Polb* deficiency does not affect Ung or TLS across AP sites in CH12-F3 cells.

**Figure 1.**
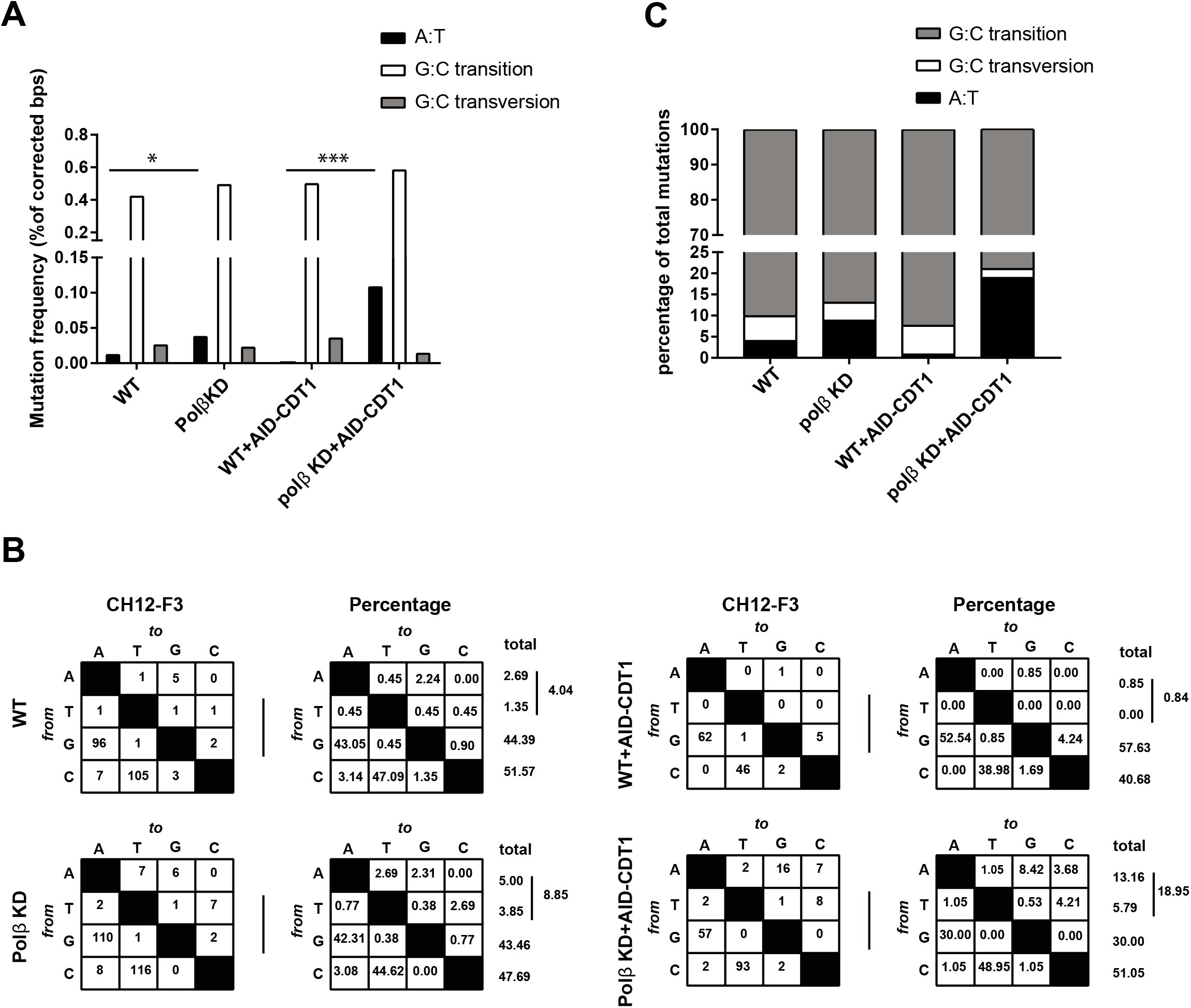
Mutations analysis in 5’Sμ sequences from WT and *Polb* knockdown CH12-F3 cells. CH12-F3 cells were stimulated with anti-CD40 antibody, IL-4 and TGF-β1(CIT) for 7 days. The 5’Sμ region was PCR amplified and mutations were analyzed by Sanger sequencing. **(A)** Mutation frequencies at A:T bps, G:C transitions, G:C transversions. Values are expressed as the percentage of total sequenced bps (corrected for the base composition). Asterisks indicate significant changes between parental and *Polb* knockdown cells (\*\**p* = 0.01, \*\*\**p*= 0.001, **** *p*= 0.00001). All the *p* values were calculated using chi-square test with Yates’ correction. **(B)** The mutation spectra of 5’ Sμ regions between parental and *Polb* knockdown cells. Values are expressed as the total number of mutations, and the percentage of total mutations from three independent experiments. The total numbers of mutations analyzed are as shown in table I. **(C)** Relative contribution of A:T mutations, G:C transitions, and G:C transversions. Values showing the percentage of total mutations.

**Table I:** SHM frequencies in the 5’Sμ region of CH12-F3 cells

|  | **WT** | **Polβ KD** | **WT+AID-CDT1** | **Polβ KD+AID-CDT1** |
| --- | --- | --- | --- | --- |
| **Total mutations** | 223 | 260 | 118 | 190 |
| **Total bp sequenced** | 106121 | 103095 | 49005 | 58092 |
| **Total mutation frequency** | $2.1 \times 10^{-3}$ | $2.5 \times 10^{-3}$ | $2.4 \times 10^{-3}$ | $3.3 \times 10^{-3}$ |
| **Mutation frequency at AID hotspots** | $4.3 \times 10^{-3}$ | $5.1 \times 10^{-3}$ | $5.4 \times 10^{-3}$ | $5.9 \times 10^{-3}$ |

### Polβ prevents A:T mutagenesis in the G1-phase of the cell cycle

Previous studies indicated that mncMMR can be activated independently of DNA replication (Peña-Diaz et al., 2012; Schrader et al., 2007). To determine whether Polβ inhibits mncMMR-mediated A:T mutations in the G1-phase of the cell cycle, we expressed AID fused to chromatin licensing and DNA replication factor 1 (Cdt1), which destabilizes protein outside of the G1-phase (Le and Maizels, 2015). It was shown that in the Ramos B-cell line, the number of A:T mutations decreased upon transduction of the cells with AID-mCherry-Cdt1 (Le and Maizels, 2015). Interestingly, expression of AID in the G1-phase in CH12-F3 resulted in a much greater increase in A:T mutations in *Polb* knockdown cells (0.4 x 10^-4^ vs 11 x 10^-4^ *p*=0.0001; 27.5-fold increase, **Fig. 1**). These data not only confirm the inhibitory effect of Polβ on the generation of A:T mutations but also shows that Polβ predominantly restricts A:T mutagenesis to the G1-phase of the cell cycle.

### Polβ prevents recruitment of MMR proteins to the 5’ Sμ region

The finding that silencing of *Polb* enabled A:T mutagenesis in CH12-F3 cells suggested that Polβ prohibits mncMMR mediated repair of AID-induced lesions. To determine whether *Polb* silencing affected the binding of MMR proteins to the 5’ side of the Sμ region (**Fig. 2A**), we carried out chromatin immunoprecipitation (ChIP) experiments in WT and Polβ knockdown CH12-F3 cells. At the 5’ Sμ region, Msh2, Msh6, and Exo1 were found enriched in *Polb* knockdown cells, confirming that Polβ prevents mncMMR downstream of AID-induced lesions (**Fig. 2B**).

**Figure 2.**
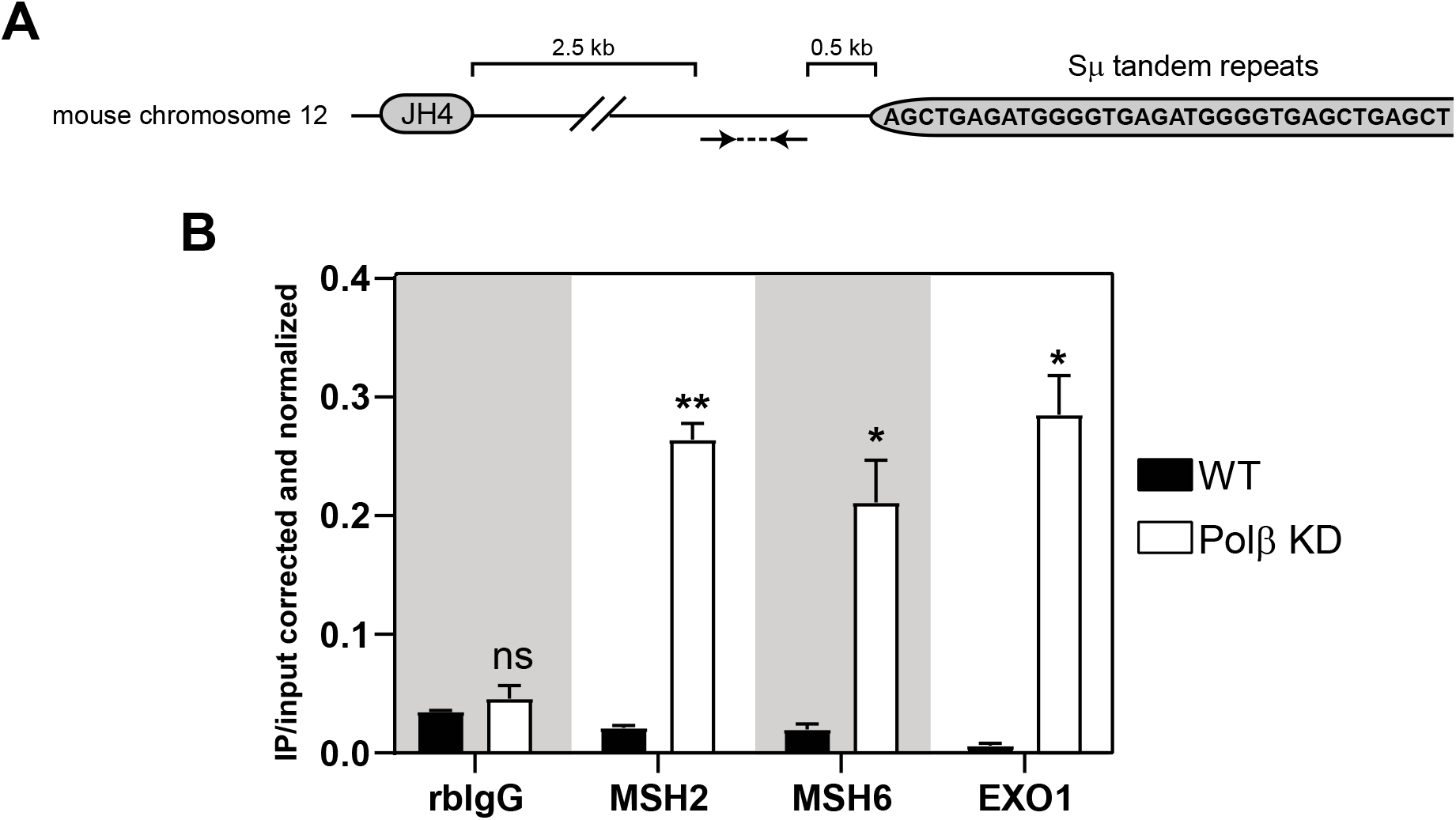
Chromatin immunoprecipitation shows enrichment of MMR components at 5’Sμ in *Polb* knockdown cells. **(A)** Schematic depiction of position of the designed primer pair (arrows) for ChIP-qPCR analysis. **(B)** Binding of polyclonal rabbit IgG (RbIgG), Msh2, Msh6 and Exo1 to the 5’Sμ region in CIT-treated CH12-F3 cells were determined by ChIP and expressed as percent of input DNA. ChIP experiments were performed at least twice. (n=3, technical replicates, *p<0.05, **p<0.01, ns = not significant, unpaired *t*-test wit Welch’s correction). Bar graphs depict means ± SEM.

### Polβ mediated inhibition of A:T mutagenesis is not restricted to B cells or Ig genes

Recently, it was shown that mncMMR is not solely active in B cells but also in other cell types (Peña-Diaz et al., 2012). In an attempt to learn how mncMMR is regulated in non-B cells, we created *Polb* knockdown mouse NIH-3T3 fibroblast cells using shRNA (**Fig. S1C**). The functional knockdown of *Polb* was confirmed by increased MMS sensitivity (**Fig. S1D**). AID fused to the estrogen receptor (AID-ER) was then overexpressed in this cell line, by which nuclear translocation of this fusion protein can be enforced by 4-hydroxy-tamoxifen (4-OHT) treatment. In addition, the retroviral overexpression construct contains a truncated nerve growth factor receptor (ΔNGFR) incapable of signaling, which allows the identification and sorting of transduced cells by staining with an anti-NGFR monoclonal antibody. In order to monitor AID activity we also transduced the cells with a fluorescent reporter plasmid (mOrange^STOP^-transgene) (Pérez-Durán et al., 2012). This reporter construct contains an amber <u>TAG</u> stop codon as part of the <u>AG</u>CT AID mutational hotspot at positions 230-233 of the sequence encoding the mOrange fluorescent protein, producing a non-fluorescent protein. However, activation of AID by 4-OHT may result in the introduction of transversions in the stop codon, reverting it to TAT or TAC, and generate full-length mOrange fluorescent protein. Moreover, the plasmid also contains GFP, which allows the tracking of transduced cells (Pérez-Durán et al., 2012). To test the mOrange^STOP^ revertance assay, WT and *Polb* knockdown NIH-3T3 cells were cultured in the presence or absence of 4-OHT for up to 11 days. mOrange+ cells appeared in AID-ER transduced cultures after 4-OHT treatment (7% mOrange+ in 4-OHT treated cells vs 0.8% in untreated, **Fig. S2**). These results confirmed that the mOrange construct is suitable to monitor AID mutational activity. Then, we analyzed mutations along the entire mOrange^STOP^ sequence from three independent experiments in WT and *Polb* knockdown NIH-3T3 cells. The cells were cultured in the presence of 4-OHT for 11 days. The mOrange^STOP^ transgene was PCR amplified from the total cell population and mutations were analyzed by cloning and Sanger sequencing. Similar to CH12-F3 B cells, *Polb* knockdown significantly increased mutation frequency at A:T bps (1.4 x10^-4^ vs 4.4 x10^-4^, *p*= 0.03, **Fig. 3**). Interestingly, in contrast to B cells we found that *Polb* knockdown also significantly increased the total mutation frequency (2.1 x10^-4^ vs 8.4 x10^-4^, *p*=0.0001) and the mutation frequency in C or G in AID hotspots (WR<u>C</u>Y/R<u>G</u>YW/WR<u>C</u>/<u>G</u>YW) (1×10^-4^ vs 5.3 x10^-4^, *p*= 0.0001, **Table II**). The difference between CH12-F3 and NIH-3T3 cells could be due to the overexpression of AID in the latter, and/or undetermined cell type-specific characteristics.

**Figure 3.**
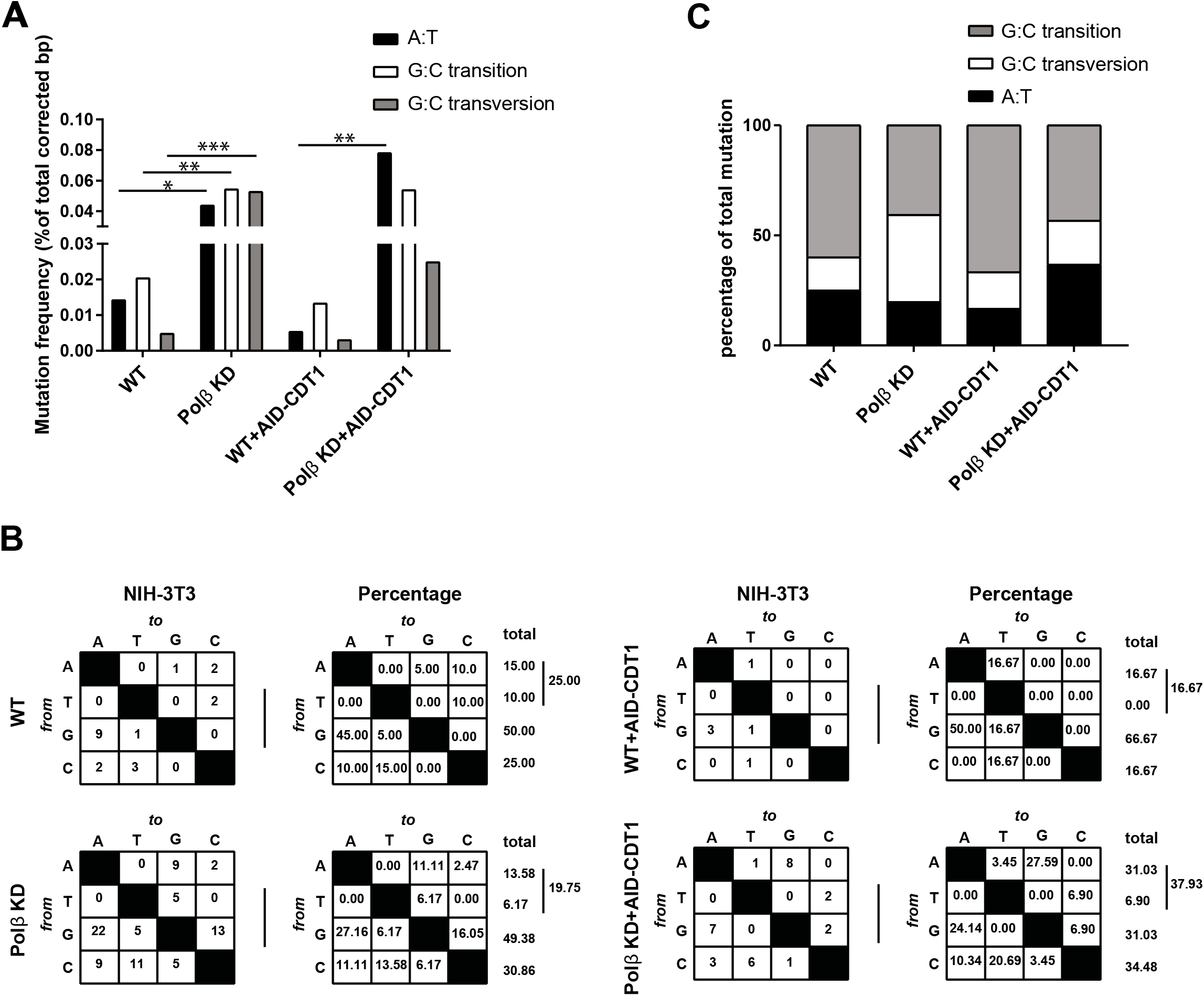
AID-induced mutation analysis in WT and *Polb* knockdown NIH-3T3 cells. NIH-3T3 cells were transduced with mOrange^STOP^-IRES-GFP and AID-ER-IRES-ΔNGFR expressing vectors. Transduced cells were cultured in the presence of 4-OHT for 11 days. mOrange+ GFP+ cells were sorted and the mOrange^STOP^ transgene was PCR amplified from the total cell population and mutations were analyzed by Sanger sequencing. **(A)** mutation frequencies at A:T bps, G:C transitions, G:C transversions. Values are expressed as the percentage of total sequenced bps (corrected for the base composition). Asterisks indicate significant changes between WT and Polβ knockdown cells (\*\**p* = 0.01, \*\*\**p*= 0.001, **** *p*= 0.00001). All the *p* values were calculated using chi-square test with Yates’ correction. **(B)** The mutation spectra of the mOrange^STOP^ transgene between WT and *Polb* knockdown cells. Values are expressed as the total number of mutations, and the percentage of total mutations from three independent experiments. The total numbers of mutations analyzed are as shown in table II. **(C)** Relative contribution of A:T mutations, G:C transitions, and G:C transversions. Values showing the percentage of total mutations.

**Table II:** SHM frequencies in the mOrange^STOP^ sequence in NIH-3T3 cells

|  | WT | Polβ KD | WT+AID-CDT1 | Polβ KD+AID-CDT1 |
| --- | --- | --- | --- | --- |
| **Total mutations** | 20 | 81 | 6 | 29 |
| **Total bp sequenced** | 94968 | 95474 | 46646 | 36665 |
| **Total mutation frequency<sup>\*\*\*</sup>** | $2.1 \times 10^{-4}$ | $8.4 \times 10^{-4}$ | $1.2 \times 10^{-4}$ | $7.9 \times 10^{-4}$ |
| **Mutation frequency at AID hotspots<sup>\*\*\*</sup>** | $1.0 \times 10^{-4}$ | $5.3 \times 10^{-4}$ | $1.0 \times 10^{-4}$ | $3.3 \times 10^{-4}$ |

Asterisks indicate significant changes between WT and Polβ knockdown cells (\*p < 0.05, \*\*p < 0.01,
\*\*\* p<0.001). All the p values were calculated using chi-square test with Yates' correction.

Interestingly, *Polb* knockdown resulted in a significant increase in the frequency of G:C transitions (2×10^-4^ vs 5.5 x10^-4^, *p*=0.004) and G:C transversions (0.5×10^-4^ vs 5.3×10^-4^, *p*=0.0001, **Fig. 3**). These data suggest that Polβ suppressed the accumulation of G:C transitions and transversions in these cells, which again confirms the faithful repair activity of this polymerase. Previously, it was demonstrated that C to G and G to C transversions can be generated by Rev1 and Polη downstream of Ung and Msh2 (Ung+Msh2 hybrid pathway) (Kano et al., 2012; Krijger et al., 2013). Interestingly, C to G and G to C transversions were significantly increased in *Polb* knockdown cells (0 vs 3.0 x10^-4^, *p*=0.0001), perhaps by preventing the formation of Msh2/6 and Exo1-dependent gaps, which are required for these transversions by Rev1 and Polη downstream of the Ung+Msh2 hybrid pathway.

In the study by Pérez-Durán *et al.* it was shown that overexpression of AID leads to an increase in the total mutation frequency and also the mutation frequency at AID hotspots (Pérez-Durán et al., 2012). We have shown that ablation of Polβ exaggerated this effect even further, which again confirms the general inhibitory effect of Polβ on AID-induced mutagenesis.

Similar to B cells we have restricted the expression of AID to G1-phase of the cell cycle using AID-mCherry-Cdt1 plasmid overexpression. The total mutation frequency and the frequency of A:T mutations did not differ significantly in WT NIH-3T3 cells expressing AID-ER versus AID-Cdt1. However, A:T mutations were increased more prominently in *Polb* knockdown cells compared to WT cells when AID-Cdt1 was expressed (from 0.6 x10^-4^ to 7.9 x10^-4^; 13.2-fold increased; WT vs *Polb* knockdown respectively) compared to cells expressing AID-ER (from 1.4 x10^-4^ to 4.4 x10^-4^; 3.1-fold increase). We found that expression of AID-Cdt1 did not further increase the total mutation frequency (**Fig. 3**). These results underscore that the Polβ-mediated inhibition of A:T mutagenesis in the G1-phase is not restricted to B cells but is also active in other cell types.

### Expression of Polβ in the Germinal Center

Strikingly, several *in vitro* systems for AID-induced mutagenesis show a relative underrepresentation of A:T mutations versus *in vivo* human or mouse GC B cells (~0-20% versus 50-60% of all mutations, respectively) (Martin et al., 2002; Matthews et al., 2014; Xiao et al., 2007; Yoshikawa et al., 2002). Based on our findings, we hypothesized that this discrepancy could be due to the difference in the regulation of Polβ under *in vitro* versus *in vivo* conditions. Therefore, we sought to determine the expression of Polβ in the GC in tissue sections of human tonsil, reactive lymph node and Peyer’s patches. Sections of human testis were used as a positive control for Polβ staining. Even though that nuclear staining of Polβ was observed in testis sections (**Fig. S3A**), we barely detected any Polβ expression in the GCs in tonsil tissue (**Fig. 4A**), reactive lymph node (**Fig. S3B**) and Peyer’s patches (**Fig. S3C**) by immunohistochemistry. Despite low expression of Polβ in the GC, we observed sporadic scattered cells with high levels of nuclear Polβ within the GC. To determine the origin of these cells we have conducted double staining for Polβ and T cells (CD3), which clearly indicated that follicular T cells are the source of Polβ expression in the GC (**Fig. 4B**). Immunofluorescence staining of Polβ confirmed the low expression in the DZ and LZ of GC as demarcated by the Ki67 proliferation marker **(Fig. 4C)**.

**Figure 4.**
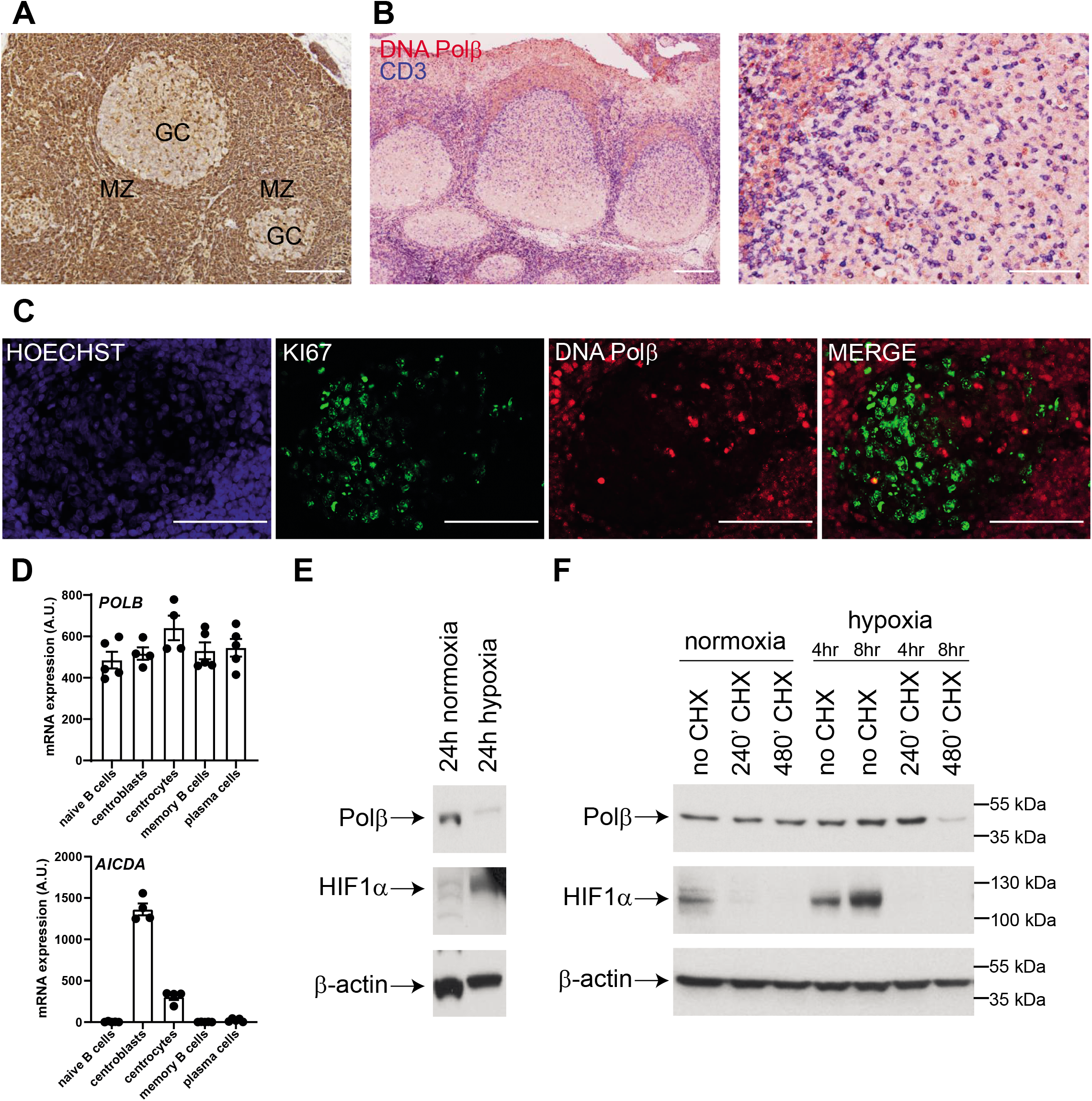
Low Polβ protein expression in germinal center B cells and hypoxia increases Polβ turnover. **(A)** Human tonsil immunohistochemical staining of Polβ (brown, nuclear staining), germinal centers (GC) and mantle zone (MZ) are indicated. Scale bar is 200 μm. **(B)** Human tonsil Immunohistochemical double staining of Polβ (in red, nuclear staining) and CD3 (= T cells; in blue, membrane staining). Scale bar is 200 μm. The germinal center consists of a dark zone, where B cells proliferate and undergo AID-dependent mutagenesis, and a light zone, where B cells interact with T cells and are selected based on antibody affinity. T follicular helper (Tfh) cells (CD3, blue membrane staining) express nuclear Polβ (red staining) (right hand panel). Scale bar is 50 μm. **(C)** Immunofluorescence images of human tonsil sections showing Hoechst33342 (blue), KI67 (green), and DNA Polβ (red). Scale bars are 200 μm. **(D)** Bar graphs depicting arbitrary units (A.U.) of normalized mRNA expression of human *POLB* and *AICDA* genes in purified peripheral blood naïve B cells (n=5), tonsillar GC centroblasts (n=4), tonsillar GC centrocytes (n=4), peripheral blood memory B cells n=5), and bone marrow plasma cells (n=5), as determined by microarray gene expression profiling (Kassambara et al., 2015). Bars represent means, error bars ± SEM, dots represent individual donors. **(E)** CH12-F3 cells were cultured for 24 h at normoxia or hypoxia in the absence of CHX. HIF1α was used as a positive control for hypoxia. **(F)** Polβ protein stability was determined by the cycloheximide (CHX) chase assay in anti-CD40 + IL4 + TGFβ (CIT) stimulated CH12-F3 cells. CH12-F3 cells were cultured with CHX for 240 minutes and 480 minutes at normoxic (~20% O_2_) or hypoxic conditions (1% O_2_), Polβ protein expression was determined by western blotting. β-actin was used as a loading control.

We compared the mRNA expression of *POLB* in sorted human naïve B cells, centroblasts, centrocytes, memory B cells and bone marrow plasma cells by analysis of a previously published gene expression profiling data set (Kassambara et al., 2015). Expression of *AICDA* was increased in the centroblasts confirming the sorting strategy and purity, whereas *POLB* mRNA was not significantly different, suggesting that Polβ expression is post-transcriptionally regulated in GC B cells **(Fig. 4D)**. Of interest, it was recently shown that the GC is a hypoxic environment (Abbott et al., 2016; Cho et al., 2016). Furthermore, in several cancer cell lines it was demonstrated that hypoxia diminished BER activity by changing the expression of several BER proteins (Chan et al., 2014). This prompted us to assess the effect of low oxygen/hypoxia conditions (1% O_2_) on Polβ protein stability in the GC B-cell-derived CH12-F3 cell line. CH12-F3 cells were stimulated with CIT for 48 h and maintained at normoxic conditions (20% O_2_) or subsequently cultured under hypoxic conditions (1% O_2_). Exposure to hypoxia for 24 h resulted in a decrease in Polβ protein expression; hypoxic conditions were confirmed by increased hypoxia-inducible factor 1 subunit alpha (HIF1α) expression (**Fig. 4E**). Next, we assessed Polβ protein stability by treating the cells with the protein synthesis inhibitor cycloheximide (CHX). Time course analysis of Polβ expression was performed by immunoblotting, showing that hypoxia accelerated Polβ protein degradation, which became apparent after 8 h of treatment (**Fig. 4F**). Interestingly, CHX treatment also resulted in the loss of HIF1α expression, suggesting that the Polβ degradation was regulated in a HIF1α independent fashion. These data suggest that A:T mutagenesis during SHM in the GC may be related to the hypoxic micro-environment, which lowers protein expression of Polβ. Strikingly, follicular T cells in the GC retain nuclear Polβ expression, whereas these cells reside in the same hypoxic environment. These observations suggest that additional (GC B-cell-specific) factor(s) may be involved in the hypoxia-triggered down modulation of Polβ. These factors remain to be identified. In contrast to our findings, Schrader *et al.* reported that Polβ protein expression was detectable in FACS-purified mouse GC B cells from Peyer’s patches (Schrader et al., 2013), but this may be related to the exposure of cells to normoxic conditions during the purification of these cells.

In summary, why in GC B cells AID-instigated dU lesions are not restored faithfully was not understood. We now demonstrate that spBER itself is generally suppressed by Polβ degradation. In this way, AID-instigated lesions are directed towards mncMMR and long patch BER (lpBER), both of which contribute to the generation of A:T mutations (Krijger et al., 2009). *Ung* deficiency had a limited effect on A:T mutagenesis (~10% reduction) (Rada et al., 2002), whereas *Msh2* or *Msh6* deficiency resulted in ~50-90% reduction of mutations at A:T bps (Rada et al., 1998; Shen et al., 2006). Interestingly, combined deficiency of *Ung/Msh2* or *Ung/Msh6* resulted in the complete ablation of A:T mutagenesis (Rada et al., 2004; Shen et al., 2006), arguing that lpBER and mncMMR contribute to A:T mutagenesis, and in line with effective recruitment of error-prone TLS polymerases, require PCNA-Ub and Polη (Krijger et al., 2009; Langerak et al., 2007).

The relative loss of Polβ in GC B cells *in vivo* in part explains how these pathways collaborate for the induction of A:T mutations, while at the same time competing for the U:G lesions. We propose that the processive nature of AID (Pham et al., 2003) provides sufficient U:G mismatches that allows the simultaneous processing by lpBER and mncMMR. The combined (focal) activity of AID at and around the *Ig* loci and the systemic break down of Polβ appears crucial for A:T mutagenesis. The mutagenic nature of DNA repair during SHM in GC B cells is directly coupled to the inactivation of spBER, one of the most efficient repair modes in mammals. We speculate that the overall loss of Polβ in combination with thousands of genome-wide deoxyuridine triphosphate (dUTP) misincorporations and spontaneous deaminations per day per cell that require spBER (Lindahl, 1993) are key contributors in destabilizing the genome of GC B cells. Our observations give insight into the regulation of SHM and explains why GC B cells should be intrinsically prone to apoptosis, not just of GC B-cell selection, but perhaps also to minimize DNA damage. The specific loss of Polβ may lie at the heart of GC-derived lymphoma development and suggests that, apart from AID activity, the shutdown of a late step in global spBER likely is an important contributor to the onset of GC B-cell lymphomas. Ongoing long-term *in vivo* experiments will define their relative contributions.

## Materials and methods

### Cell culture

Mouse CH12-F3 lymphoma cells (a kind gift from Prof. Tasuku Honjo, Kyoto University, Japan) were maintained in RPMI 1640 (Life Technologies, Bleiswijk, The Netherlands) supplemented with 10% FCS, 50 μM 2-mercaptoethanol and 5% NCTC-109 medium (Gibco, Waltham, MA). For induction of SHM, the cells were stimulated with 1 μg/ml anti-CD40 antibody (clone HM40-3; eBioscience, San Diego, CA), 5 ng/ml mouse IL-4 (ProSpec, Rehovot, Israel), and 0.5 ng/ml mouse TGF-β1 (ProSpec), and cells were grown for 7 days. Mouse NIH-3T3 fibroblast cells were obtained from (ATCC^®^ CRL-1658™, Manassas, VA) cultured in DMEM (Life Technologies), supplemented with 2 mM of L-glutamine, 100 U/ml penicillin, 100 mg/ml streptomycin. Cells were treated with 1 μM 4-OHT (Sigma Aldrich, St. Louis, MO) where indicated.

### Retroviral transduction

Retroviral particles were generated by transient transfection of HEK293 cells with pCL-ECO (Imgenex, Cambridge, UK) packaging construct in combination with pSuper.mBeta826 (DNA Polβ shRNA, a kind gift from Dr. Robert Sobol, [Addgene, Plasmid #12549]) (Trivedi et al., 2005) or AID-ER (a kind gift from Dr. Vasco Barretto, instituto Gulbenkian de Ciência, Oeiras, Portugal) and pMX-PIE-Orange^STOP^ (a kind gift from Dr. Almudena Ramiro, CNIC, Madrid, Spain) (Pérez-Durán et al., 2012) and AID-mCherry-Cdt1 (a kind gift from Nancy Maizels, University of Washington School of Medicine, Seattle, WA) (Le and Maizels, 2015) retroviral vectors using FuGene (Promega, Madison, WI) transfection reagent. Forty-eight hours after transfection, retrovirus-containing supernatant was collected and passed through 0.45 μm filters.

The retrovirus was first bound to the plate coated with RetroNectin (Takara, Kusatsu, Japan) and centrifuged for 3 h at 2,000x g at 4°C and cells were added after removing the retrovirus supernatant. For the generation of polβ knockdown cells, transduced cells were selected by puromycin treatment. Whole cell extracts were prepared and analyzed by western blotting. Parental wild type (WT) control cells retained expression of Polβ, while the shRNA-transduced cells had nearly undetectable levels of Polβ. Expression of DNA Polβ was checked throughout the course of the experiments. After the transduction of NIH-3T3 cells with AID-ER and pMX-PIE-Orange^STOP^, cells were cultured in the presence of 1 μM 4-OHT, and GFP+ mOrange+ cells were monitored and sorted by flow cytometry (FACSCanto; BD Biosciences, San Jose, CA) at the indicated time points.

### Cytotoxicity studies by growth inhibition assay

CH12-F3 sells were seeded (20×10^4^ per well) in 24 well plate in 1ml medium in triplicate without selection antibiotics. The next day, cells were treated for 1 h with a range of concentrations of the DNA methylating agent methyl methanesulfonate (MMS) (Sigma-Aldrich), and subsequently cultured for an additional 48 h. Cell death was assessed by 7-aminoactinomycin D (7-AAD, Affymetrix eBioscience, San Diego, CA) staining and flow-cytometry (FACSCantoII^™^, BD Biosciences). The percentage of dead cells was normalized to the untreated condition. For the NIH-3T3 cell line, cells were exposed to MMS for 24 h. Viability was assessed using the MTT (3-(4,5-dimethylthiazol-2-yl)-2,5-diphenyl tetrazolium bromide tetrazolium; Sigma Aldrich) assay, data was acquired using a CLARIOstar^®^ Plus plate reader (Isogen Life Science, De Meern, The Netherlands). Data was normalized to the untreated condition.

### Chromatin immunoprecipitation

Protein-DNA binding was studied using the MAGnify Chromatin Immunoprecipitation system (Life Technologies) according to the manufacturer’s protocol. In brief, 10×10^6^ cells were used for each IP, which were performed in triplicate for each condition. The cells were fixed in 1% formaldehyde for 5 minutes at 37°C and quenched for 10 min with 125 mM of glycine. Subsequently, cells were washed twice in ice-cold PBS and lysed using the enclosed lysis buffer and sonicated by Covaris S2 (intensity: 3, duty cycle: 5%, and cycles per burst: 200) to obtain DNA fragments of an average length of 500 bp. Protein concentrations were measured after sonication by the bicinchoninic acid assay (Bio-Rad, Hercules, CA), and equal amounts of fragmented chromatin–protein complexes were incubated for 2 h with enclosed Dynabeads coupled to rabbit anti-Polβ (Ab26343, Abcam, Cambridge, UK), mouse anti-MSH2 mAb (FE11, Calbiochem, San Diego, CA); mouse anti-MSH6 mAb (610919, BD biosciences); and rabbit anti-EXO1 pAb (NBP1-19709, Novusbio, Cambridge, UK) (10 μg of each Ab per ChIP); or to polyclonal rabbit IgG (control ChIP) (Life Technologies). After a series of washes, chromatin was de-crosslinked for 15 minutes at 65°C and purified using the ChIP DNA Clean & Concentrator kit (Zymo Research, Freiburg, Germany). Binding of Msh2, Msh6, polβ and Exo1 to the 5’Sμ (in CH12-F3) was assessed by quantifying ChIP-enriched DNA using CFX384 realtime PCR (Bio-Rad). The following primers were used for qPCR:

5’Sμ-Forward: 5’-TCTGTACAGCTGTGGCCTTC-3’

5’Sμ-Reverse: 5’-GATCCGAGGTGAGTGTGAGA-3’

### Western blot analysis

Cells were washed with ice-cold PBS and lysed in ice-cold lysis buffer (10 mM Tris-HCl pH 8, 140 mM of NaCl, 1% Nonidet P-40, 0.1% sodium deoxycholate, 0.1% SDS, 1 mM of EDTA) supplemented with protease and phosphatase inhibitors (EDTA-free protease mixture inhibitor; Roche Diagnostics, Almere, the Netherlands). The protein pellets were homogenized passing through a 27G needle, and protein concentrations were measured using the bicinchoninic acid assay. For each sample 10 μg of total protein lysate was used for protein separation in Precise 4–20% gradient Tris-SDS gels (Bolt, Thermo Fisher Scientific, Landsmeer, The Netherlands). Subsequently, separated protein lysates were transferred onto polyvinylidene difluoride membranes (Immobilon-P; EMD Millipore, Burlington, MA); membranes were blocked in 5% BSA (BSA Fraction V; Roche Life Sciences, Almere, The Netherlands) in TBS-T or 5% milk in TBS-T for 1 h. Primary Abs were incubated overnight at 4°C. After a series of thorough washes with TBS-T, membranes were incubated with secondary Abs (goat anti-rabbit HRP or rabbit anti-mouse HRP; DAKO, Agilent Technologies, Heverlee, Belgium) for 2 h at room temperature. Ab binding (protein expression) was visualized using Amersham ECL Prime Western blotting detection reagent (GE Healthcare Bio-Sciences AB, Uppsala, Sweden). The following Abs were used in this study: mouse anti-Polβ (18S, Thermo Fisher Scientific); rabbit anti-HIF1-α (NB100-449, Novusbio) and mouse anti-β-tubulin (ab15568, Abcam). In the experiments using the protein translation inhibitor cycloheximide (Sigma Aldrich), CH12-F3 cells were exposed to low oxygen conditions (1% O_2_) using a Whitley hypoxic workstation (Whitley, Bingley, UK) for up to 8 h or cultured at normoxic conditions (20% O_2_).

### Immunohistochemistry and immunofluorescence staining

For immunohistochemical analysis, the slides of human tonsil, testis, reactive lymph node and Peyer’s patch tissue were obtained from the Amsterdam University Medical Center (location AMC, Amsterdam, The Netherlands) with informed consent. This study was conducted in accordance with the ethical standards in our institutional medical committee on human experiments, as well as in agreement with the Helsinki declaration of 1975, revised in 1983. Tissue sections were deparaffinized with xylene and hydrated through an alcohol rinse series follow by 15 minutes incubation in methanol + 0.3% H_2_O_2_. After heat inducible epitope retrieval in citrate buffer (pH=6) in a pressure cooker for 20 minutes, slides were treated with Super Block (AAA125, ScyTek Laboratories, West Logan, UT) for 15 minutes at room temperature. Then, the tissue sections were incubated with rabbit anti-Polβ antibody (ab26343, Abcam). Antibody–antigen detection was performed with goat anti-rabbit poly horse radish peroxidase (HRP) (Bright Vision Poly HRP-Anti Rabbit IgG, Immunologic, VWR, Duiven, The Netherlands). The end products were visualized with 3,3’ Diaminobenzidine (DAB) (Bright DAB substrate kit, Immunologic).

For double immunohistochemistry stainings, similar to single staining, human tonsil tissue sections were deparaffinized. After heat induced-epitope retrieval and blocking with Super Block, T cells were stained with rabbit anti-CD3 (ab16669, Abcam) overnight at 4°C. After several washing, slides were incubated with secondary goat-anti-rabbit-Poly-alkaline phosphatase (AP) (Bright Vision Poly AP-anti Rabbit IgG, Immunologic) for 30 minutes at room temperature, followed by adding VECTOR^®^ Blue AP substrate, which resulted in a blue precipitate. The slides were then boiled in citrate buffer (pH=6) again to remove the bound Abs, follow by blocking with Super Block and restained with rabbit anti-Polβ (ab26343, Abcam), followed by goat anti-rabbit-Poly-AP (Bright Vision Poly AP-anti Rabbit IgG, Immunologic) for 30 minutes at room temperature and subsequently incubated with VECTOR^®^ Red AP substrate, resulting in a red precipitate. The slides were covered with VECTAMOUNT^®^ Permanent Mounting Medium (H-5000, Vector laboratories, Burlingame, CA).

For immunofluorescence stainings, human tonsil tissue sections were deparaffinized and subjected to antigen retrieval as described above. After blocking and permeabilization in 1× PBS + 0.01% Tween-20 + 0.1% BSA, sections were stained with rabbit anti-Polβ (Ab26343, Abcam) and mouse anti-Ki67 (MIB-1, Agilent DAKO, Santa Clara, CA) overnight at 4°C, followed by goat anti-mouse IgG H&L Alexa Fluor^®^ 488 (Ab150113, Abcam) and polyclonal donkey anti-rabbit IgG Alexa Fluor^®^ 594 (711586152, Jackson Immuno Research Labs, Ely, UK). Nuclei were counterstained with Hoechst33342 (Sigma Aldrich) and slides were covered with VECTASHIELD^®^ antifade mounting medium (Vector Laboratories). Immunofluorescent images were captured using a Leica DM5500B fluorescence microscope and the Leica LAS X software suite (Leica, Wetzlar, Germany).

### mOrange^STOP^ amplification and mutation analysis

For analysis of mutations at the mOrange^STOP^ transgene, co-transduced NIH-3T3 cells were cultured for 11 days in 6 separate wells in the presence of 1μM 4-OHT. Cells were harvested, pooled and GFP+ and mOrange+ cells were sorted by flow cytometry. DNA was extracted using QIAamp DNA mini kit (Qiagen, Hilden, Germany) and mOrange^STOP^ was amplified using the following primers:

(forward) 5’-AATAACATGGCCATCATCAAGGA-3’

(reverse) 5’-ACGTAGCGGCCCTCGAGTCTCTC-3’

using Phusion High-Fidelity DNA Polymerase kit (Thermo Fisher Scientific) in five independent 25 μl PCR reactions, under the following conditions: 98°C for 30s, followed by 35 cycles at 98°C for 10 s, 65°C for 30s, and 72°C for 45s.

For Sanger sequencing, PCR reactions were pooled and products were cloned using the Zero Blunt PCR cloning kit (Invitrogen) and plasmids from individual colonies were sequenced using M13 universal primers (forward) 5’-GTAAAACGACGGCCAG-3’ and (reverse) 5’-CAGGAAACAGCTATGAC-3’. Sequence analysis was performed using Codon Code Aligner software (Codon Code Corp., Centerville, MA).

### 5’Sμ amplification and mutation analysis

For mutation analysis at the Sμ region, CH12-F3 cells were stimulated with 1 μg/ml anti-CD40 antibody (HM40-3, eBioscience), 5 ng/ml mouse IL-4 (ProSpec), and 0.5 ng/ml mouse TGF-β1 (ProSpec), and cells were grown for 7 days in 6 separate wells. DNA was extracted and the 5’Sμ fragment was PCR amplified using the primers that were reported previously (Matthews et al., 2014). 50ng of DNA was amplified in five independent PCR amplification reactions, which were pooled and sequenced, and analyzed as described for the mOrange^STOP^ gene. Cloned 5’ Sμ fragments were aligned with germline Sμ sequenced from C57BL/6 chromosome 12 (available from GenBank/EMBL/DDBJ under accession number AC073553).

### Statistics

For Sanger sequencing experiments, *p* values were calculated by Chi-square with Yates’ correction. For ChIP experiments means ± SEMs are represented. Comparisons between groups were analyzed using unpaired *t* test with Welch’s correction using the GraphPad Prism software package (GraphPad Software, La Jolla, CA).

## ACKNOWLEDGEMENTS

This study is part of a project (COSMIC; www.cosmic-h2020.eu) that has received funding from the European Union’s Horizon 2020 research and innovation program under the Marie Skłodowska-Curie grant agreement No. 765158. This research was supported by the Netherlands Organization for Scientific Research Innovational Research Incentives Scheme VIDI grant no.16126355, and an AMC Fellowship grant (both to J.E.J.G.), and the NWO/ZonMW Top grant 91213018 to H.J.

## AUTHORSHIP AND CONFLICT-OF-INTEREST STATEMENTS

J.E.J.G. and MB designed the research; M.B., B.P., M.S., T.P.vD. S.M. performed the experiments; J.E.J.G., M.B, C.v.N. and H.J. analyzed the data; J.E.J.G., M.B. wrote the manuscript; and all authors edited the manuscript.

