## Supplemental Figure 1 for "DNA polymerase β prevents AID-instigated mutagenic non-canonical mismatch DNA repair"

### Figure S1

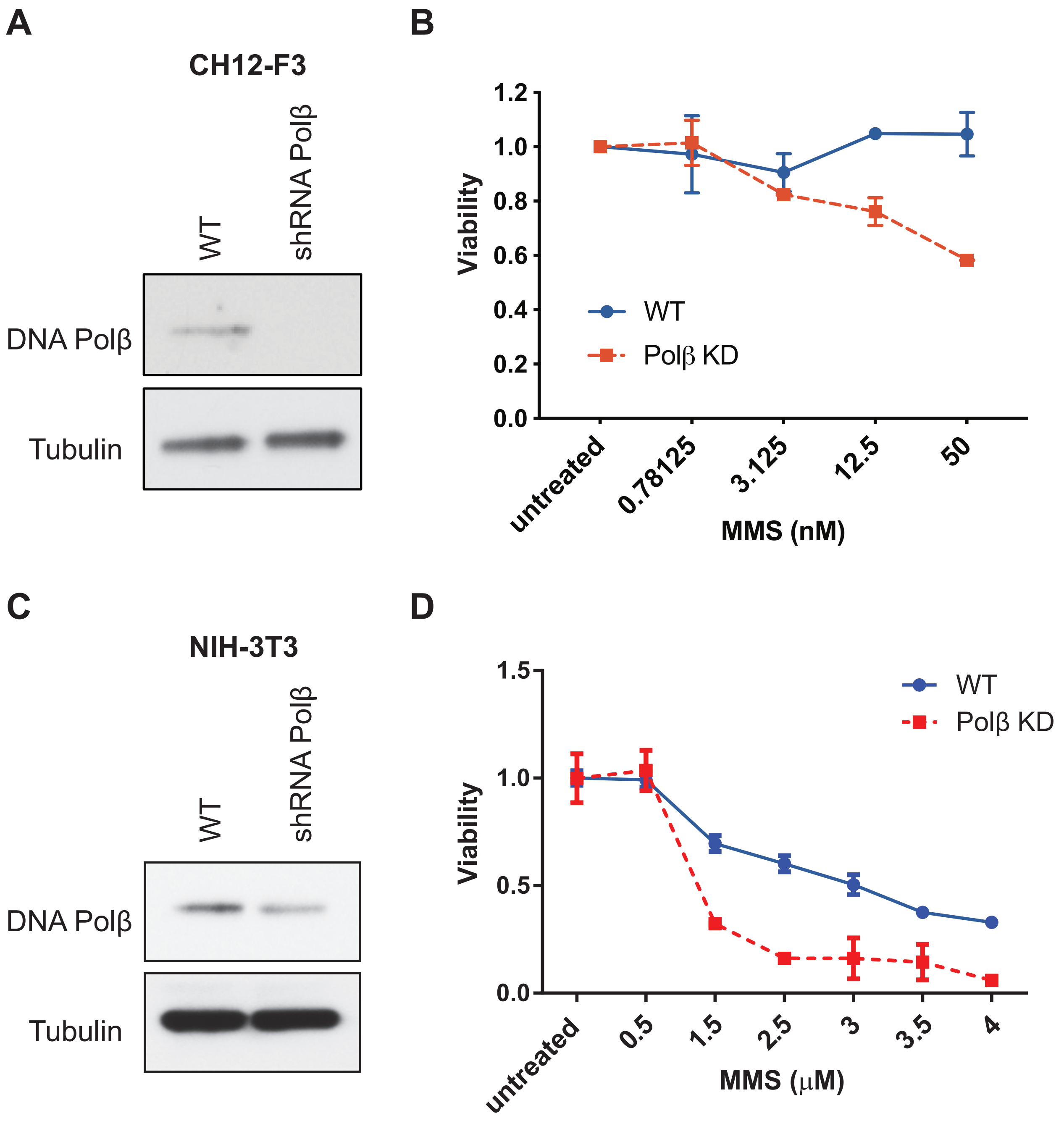


**Figure S1. Functional confirmation Polb knockdown in CH12-F3 and NIH-3T3 cells.** Immunoblot analysis showing depletion of Polβ by short hairpin-mediated silencing in CH12-F3 **(A)** and NIH-3T3 **(C)**. Tubulin was used as a loading control. WT and Polβ knockdown cells were exposed to MMS at indicated doses for 1h (CH12-F3) **(B)** or 24h (NIH-3T3) **(D)**. Graphs depict viability of cells as assessed by 7-AAD flow-cytometry (CH12-F3) or MTT assay. Three independent experiments were performed, cultures were done in duplicate, means +/- SEMs are shown.
