## Supplemental Figure 2 for "DNA polymerase β prevents AID-instigated mutagenic non-canonical mismatch DNA repair"

### Figure S2

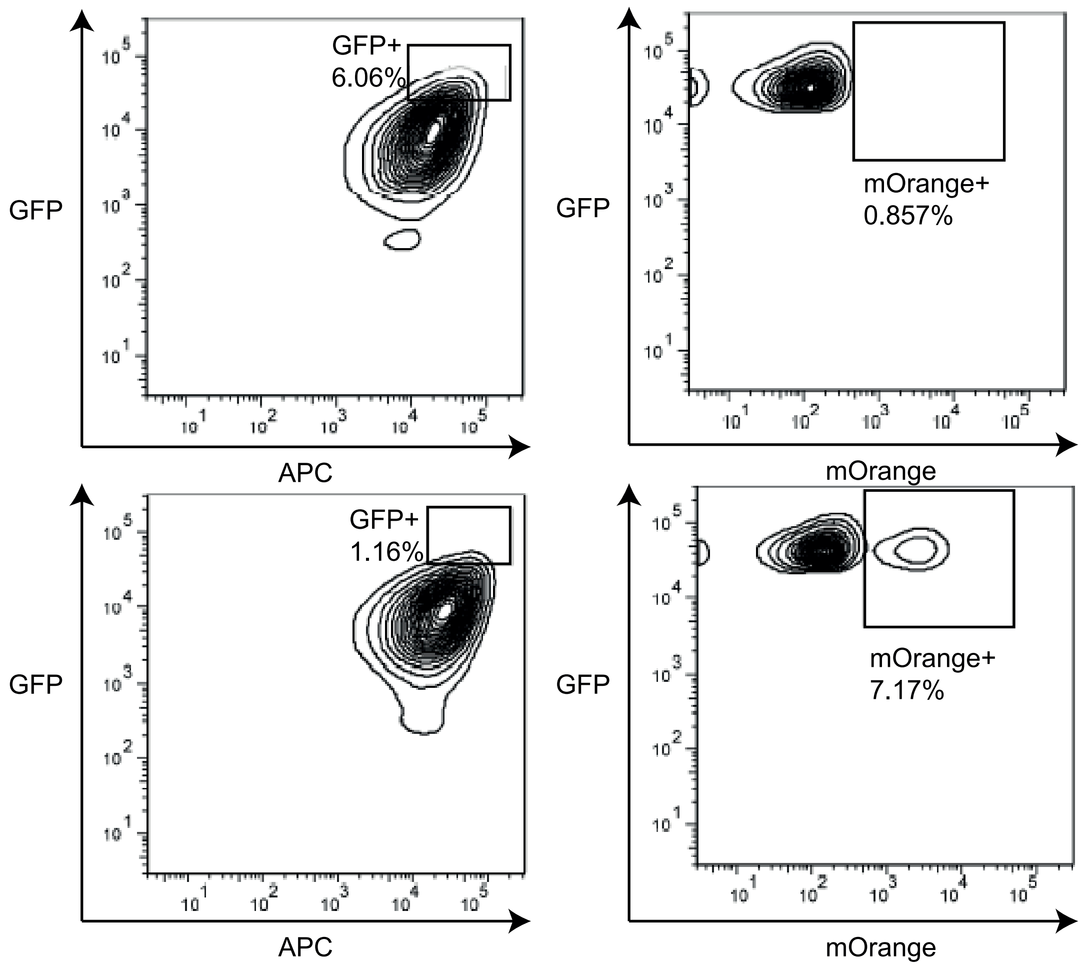


**Figure S2. mOrange$^{STOP}$ revertance assay in NIH-3T3 cells expressing AID-ER.** NIH-3T3 cells transduced with mOrange$^{STOP}$-IRES-GFP and AID-ER-IRES-ΔNGFR were stained with anti-NGFR-APC (Miltenyi Biotec, Bergisch Gladbach, Germany). Percentage of mOrange+ cells were determined after 11d of 4-OHT treatment. Percent of mOrange+ cells were determined by gating on GFP++ APC++ cells. Percentages of positive cells are indicated in the FACS plots. Upper panels show control cultures without 4-OHT, lower panels shows cultures treated with 4-OHT.
