## Supplemental Figure 3 for "DNA polymerase β prevents AID-instigated mutagenic non-canonical mismatch DNA repair"

### Figure S3

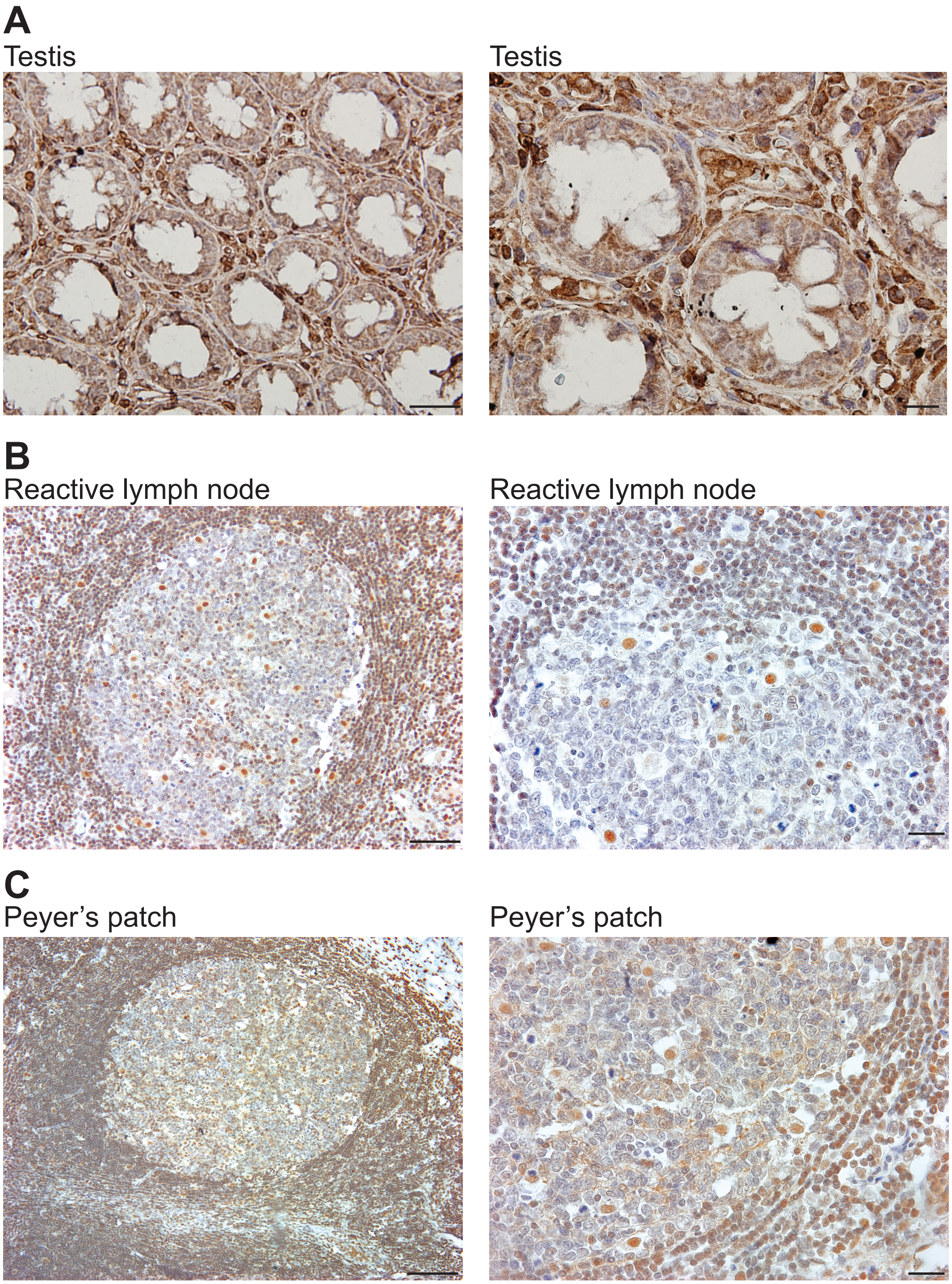


**Figure S3. Polb immunohistochemistry of human tissue specimens.** Immunohistochemical staining with anti-PolB antibody of paraffin-embedded histological sections from different human tissues. **(A)**. testis. **(B)**. Reactive lymph node. **(C)**. Peyer’s patch. Magnifications are 10x (lefthand panels; scale bar 0.05 mm) and 40x (righthand panels; scale bar 0.01 mm).
